# Biopipe: A Lightweight System Enabling Comparison of Bioinformatics Tools and Workflows

**DOI:** 10.1101/201186

**Authors:** Saima Sultana Tithi, Jiyoung Lee, Liqing Zhang, Song Li, Na Meng

## Abstract

Analyzing next generation sequencing data always requires researchers to install many tools, prepare input data compliant to the required data format, and execute the tools in specific orders. Such tool installation and workflow execution process is tedious and error-prone, and becomes very challenging when researchers need to compare multiple alternative tool chains. To mitigate this problem, we developed a new lightweight and portable system, Biopipe, to simplify the creation and execution of bioinformatics tools and workflows, and to further enable the comparison between alternative tools or workflows. Biopipe allows users to create and edit workflows with user-friendly web interfaces, and automates tool installation as well as workflow synthesis by downloading and executing predefined Docker images. With Biopipe, biologists can easily experiment with and compare different bioinformatics tools and workflows without much computer science knowledge. There are mainly two parts in Biopipe: a web application and a standalone Java application. They are freely available at http://bench.cs.vt.edu:8282/Biopipe-Workflow-Editor-0.0.1/index.xhtml and https://code.vt.edu/saima5/Biopipe-Run-Workflow

**Supplementary information:** Supplementary data are available online.

## 1 Introduction

The analysis of genome sequencing data usually requires creating multistep workflows. For example, the SNP calling from DNA sequencing data consists of two main steps: short read alignment and variant calling. Many tools are available for each of these steps e.g. the OMICTools (Henry, et al., 2014) website listed 57 different tools for the alignment step. As different tools require different platforms and library dependencies, manually installing and executing all tools for a multi-step workflow are tedious and error-prone. Things become even worse when investigate different combinations between the tools across steps to identify the best tool chain for a workflow implementation.

Many systems have been developed to create and execute workflows. Examples include Galaxy (Goecks, et al., 2010), Taverna (Hull, et al., 2006), Bioboxes (Belmann, et al., 2015), systemPipeR (Backman and Girke, 2016), ExScalibur (Bao, et al., 2015). However, none of these systems support simple execution of alternative tool chains or comparison of the performance of alternative workflows. For example, systemPipeR is an extensible environment for building and running end-to-end analysis workflows. ExScalibur allows users to easily compose DNA sequencing analysis workflows. Nevertheless, both systemPipeR and ExScalibur require users to install all tools manually. Galaxy enables users to draw a tool chain in a web browser, and then automatically installs and executes this tools chain. However, it does not facilitate users to specify alternative tools for each step in a workflow; neither does it enumerate all possible combinations between the tools across steps. If a user wants to use Galaxy as a standalone application on a local machine, the full Galaxy system must be installed which is unnecessary and is wasting a lot of computing resources.

**Figure 1:**
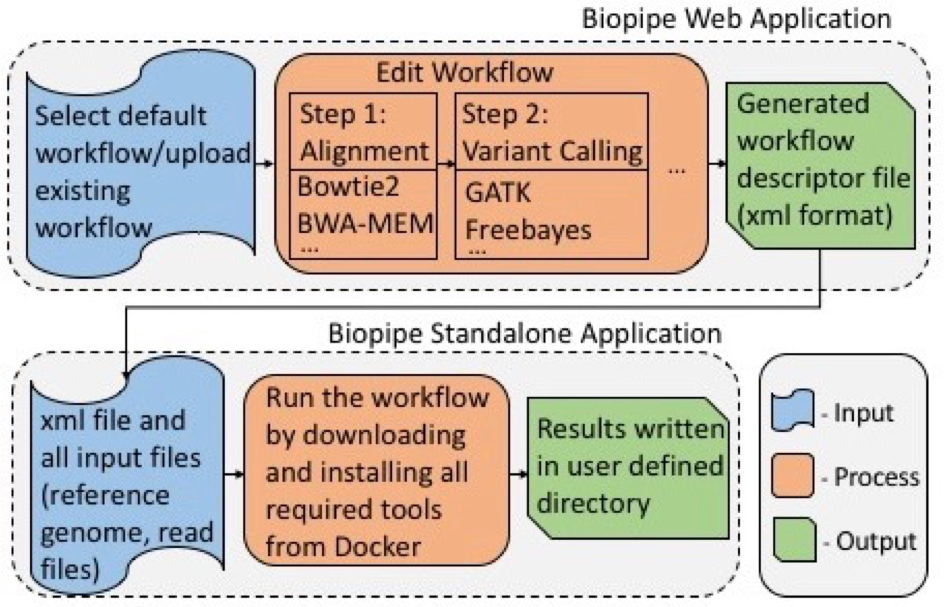
Overview of system architecture and data flow of Biopipe

To support users to specify and execute multiple alternative tool chains for a workflow, we developed a new, lightweight system, Biopipe, which leverages Docker (Merkel, 2014) ---a container-based virtualization technique---to significantly reduce the repetitive manual effort of installing and configuring tools. Biopipe is deployed as a web application together with a standalone Java application. The web application provides an easy-to-use interface for users to create and edit a workflow by specifying all alternative tools for each step. Specifically, through selecting the type of workflow and choosing one or more tools for each step, users can define numerous tool chains. Biopipe searches DockerHub (https://hub.docker.com) for Docker images that contain executables of user selected tools. Biopipe then creates a workflow file and the users can execute the workflow in any machine with Biopipe’s standalone Java application by specifying necessary input and output files, without manually installing or invoking any tool.

Biopipe’s standalone Java application automatically downloads all relevant Docker images, reuses the tool installation in each image, and synthesizes alternative tool chains by generating different Docker image compose files (compose.yml). Each compose file corresponds to a workflow step, defining how to execute the alternative tools in this step with the various data output by a previous step. Biopipe is highly portable, can be executed on Linux, Windows, Mac OS, or on the cloud. Additionally, Biopipe is lightweight for two reasons. First, a Docker image is lightweight and Biopipe only downloads and executes the minimum necessary tools selected by the users. Second, Biopipe executes every selected tool exactly once for each unique data input to eliminate repetitive computation or duplicated data storage.

## 2 Methods

Biopipe has two components, a web application and a standalone application (Figure 1). The web application of Biopipe is available at http://bench.cs.vt.edu:8282/Biopipe-Workflow-Editor-0.0.1/index.xhtml. Currently, two types of workflows are supported, ‘SNP/Indel Calling’ and ‘RNA-Seq’ workflow. The ‘SNP/Indel Calling’ workflow contains two steps: alignment and variant calling. The ‘RNA-Seq’ workflow contains two steps: read mapping and assembly. After selecting the tools for each step of a workflow, users can download a workflow descriptor file (an xml file). The file contains the name of the tools, the Docker image name for the selected tools, and the execution order of the tools. In addition to selecting a given workflow, user can also upload an xml file previously generated by our web application or a Galaxy workflow file (.ga file). After uploading such files, users can modify them by changing the selected tools for each step and then download the updated workflow files.

The standalone application of Biopipe is a platform-independent, Java GUI program and is available at https://code.vt.edu/saima5/Biopipe-Run-Workflow. Users only need to install Java and Docker prior to running this application. User can execute the same workflow file with different input data sets. After running all the tools, output will be saved in a directory selected by the user. A new tool can be added to Biopipe by developing a Docker image and a running command (in bash script) for the tool.

Biopipe was tested by creating a snp/indel calling workflow containing two alignment tools (BWA-MEM, Bowtie2) and four variant calling tools (GATK-UnifiedGenotyper, GATK-HaplotypeCaller, Freebayes, Platypus). It was also tested by creating a RNA-seq workflow containing two RNA-seq alignment tools (STAR, Tophat2) and three RNA-seq assembly tools (StringTie, Cufflinks, Trinity). A detailed description of the experiment setup and results are given in the supplementary file.

## 3 Conclusion

We developed Biopipe, a new Docker-based system to assist the researchers in creating and executing workflows in a user friendly and efficient manner. Whenever researchers need to compare and evaluate some alternative tools or alternative workflows, they need to go through the same installation and execution process, and the same steps for making the input data compatible with the tool requirement repeatedly. Biopipe can relieve the researchers from such time-consuming and error-prone processes. For future work, we will extend Biopipe by enhancing the implemented workflows with more features such as adding data preprocessing and post-processing steps, evaluating the results and generating reports, and incorporating additional alternative tools. We expect Biopipe to be of great help to the research community in the matter of executing and comparing alternative bioinformatics tools and workflows.

## Funding

This work has been supported by the Virginia Agricultural Experiment Station (Blacksburg) and the USDA National Institute of Food and Agriculture to SL.

### Conflict of Interest

none declared.

## References

Backman, T.W. and Girke, T. systemPipeR: NGS workflow and report generation environment. BMC bioinformatics 2016;17(1):388.

Bao, R., et al. ExScalibur: A high-performance cloud-enabled suite for whole exome germline and somatic mutation identification. PloS one 2015;10(8):e0135800.

Belmann, P., et al. Bioboxes: standardised containers for interchangeable bioinformatics software. Gigascience 2015;4(1):47.

Goecks, J., Nekrutenko, A. and Taylor, J. Galaxy: a comprehensive approach for supporting accessible, reproducible, and transparent computational research in the life sciences. Genome biology 2010; 11(8):R86.

Henry, V.J., et al. OMICtools: an informative directory for multi-omic data analysis. Database 2014;2014.

Hull, D., et al. Taverna: a tool for building and running workflows of services. Nucleic acids research 2006;34(suppl_2):W729–W732.

Merkel, D. Docker: lightweight linux containers for consistent development and deployment. Linux Journal 2014;2014(239):2.

